# Structure simulation of thrombopoietin receptor C-MPL with missense SNPs and its geographical distribution

**DOI:** 10.1101/342089

**Authors:** Weizhong Zheng, Dongyu Zeng, Qi Zhou, Zhilan Li, Haojian Liang

## Abstract

Mpl is a key gene controlling the process of proliferation and differentiation of megakaryocyte and several studies reported that the mutation of mpl will even cause accurate megakaryocyte leukemia. By overlapping the missense mutation in NCBI and recorded SNP in 1000 genome database, we sorted out 4 SNPs(rs190983971, rs563996763, rs546510242, rs373621350) to find out its conformational changes. With Phyre v2.0, we simulated the secondary and tertiary structure change caused by the SNP and compared it with the original structure. The significant changes indicated its potential in leading to diseases associated with platelet reproduction. Finally, according to 1000 genome database, we constructed the geographically distribution of different populations and its SNPs carrying rate.

## I. INTRODUCTION

The diversity among individuals is somewhat the diversity of their gene. DNA marker can directly manifest the differences of DNA molecular. Its high diversity, large quantity and the property that targets are either Dominant or co-Dominant state makes DNA marker a better signature than the traditional marker, including morphological marker, cellular marker, biochemistry marker. SNP (single nucleotide polymorphisms), as a new and highly accepted DNA marker, is the third generation molecular labeling technology developed from the second generation like SSR and ISSR.

SNP is widely used in many studies today not only for its high stability and rich target sites in cells but also for its wide distribution, high representation, dimorphism and easily detected. Based on the subsequent influence on genetic morphology, SNPs can be further subdivided into protein coding SNP and non-protein-coding SNP depending on whether it is located in the transcription region. Among protein coding SNPs, if the SNP can cause changes in the amino acid sequence, it will be called synonymy coding SNP. If not, then the SNP is missense coding SNP which often affect the protein function[1-3].

Leukemia is kind of cancer which is caused by the abnormal malignance proliferation of leukocyte in bone marrow driven by the cellular DNA mutation. TPO receptor gene mpl,locate in the first chromosome 1p34, contained 12 exons and transcript protein as long as 635 ~ 680 amino acid. Mpl, also called thrombopoietin receptor, is one of the most important gene controlling the proliferation and progression of megakaryocyte. Most Myeloproliferative Neoplasm (MPN) patients are diagnosed with mpl mutation, mostly seen at W515L, which suggest a high sensitivity to TPO in marrow cells and activate the non-cytosine dependent JAK-STAT pathway, resulting in the cellular malignant replication and leading to MPN tumorigenesis. Pardanani et al has also found mpl W515L mutation in Primary Myelofibrosis (PMF) and Primary Thrombocytosis (ET) while no mutation in this site have been reported in PV patients. 3% of the ET patients and 10% of the PMF patients carried W515L, whereas this ratio rise to 15% in the JAK2 V617F -/- patients[1, 4]. Beer et al also discovered that compared with JAK2 V617F -/- patients, mpl W515L carriers are in the older age and have a high risk of Arterial Thrombosis (p < 0.01). However, if compared with JAK2 V617F +/+ patients, mpl W515L carriers show a significant lower hemoglobin level but higher platelets counts (p< 0.01)[5]. Their study suggested that the W515L mutation in mpl doesn’t interference the life period of MPN patients and doesn’t deteriorate the Myelofibrosis progress, leukemia transference progress or other complication[6].

According to the SNP database affiliated to NCBI, collected data show 295 missense SNPs reported in Homo sapiens mpl gene. 1000 genome database only collected 100 total SNPs (including synonymy coding SNP, missense SNP and stop-coding SNP). By filtrating the superposition data of these two database, we can find out whether there are SNP that can probably bring amino acid shift in protein sequence. Among them, 4 SNPs are filtrated out (SNP ID:rs190983971, rs563996763, rs546510242, rs373621350). By analyzing these 4 missense SNPs, we can roughly draw a picture of the carrying rate of mpl mutation in different region and different population globally, which can indicate the risk of Megakaryocyte Associated diseases like AML, ET and PMF.

## II. METHOD

### Screening of diseases related SNPs

To sort out SNPs related to Megakaryocyte diseases, we carried out an overlap analysis of the NCBI database and 1000 genome database and finally, we found out 4 missense SNPs but no stop-code SNP. The ID of the 4 missense SNPs are rs190983971, rs563996763, rs546510242,rs373621350.

### Protein Structure Simulation

To further investigate the four mutations, we ran the Phyre v2.0 tool (http://www.sbg.bio.ic.ac.uk/phyre2/html/page.cgi?id=index) to predict the conformational change. This tool adapted the profile-profile matching algorithm to predict the protein 3D structure. We first use the origin sequences in the Intensive module in Phyre to match reported cytosine-receptor structure. The simulation was viewed in the 3D structure viewer Chimera v1.11.2 developed by USCD.

### SNP distribution

Among the 26 sub-population branches collected in the 1000 genome database, population and region without overlapped SNPs were labeled with light blue. In the region where the 4 SNPs spread (rs190983971, rs563996763, rs546510242, rs373621350), we labeled with red, green, yellow and purple. However, due to the synchronously distribution of 2 SNPs in Pakistan and Bangladesh, we chose orange to labeled rs190983971 and rs546150242. Since rs373621350 only spread in Bangladesh, the continue inheritance of purple would not cause confusion. The global map was downloaded from http://www.wpmag.org.

## III. CONCLUSION

### Missense SNPs change protein structure in simulation

To investigate the protein conformational shift affected by the 4 SNPs (Figure 1), we edited the protein sequence according to the mutation site and replaced amino acid. The altered sequence was paired with the wild type sequence and then we sent them as input uploaded to the Phyre website. Return structures were shown in Figure 2.

**Figure 1.**
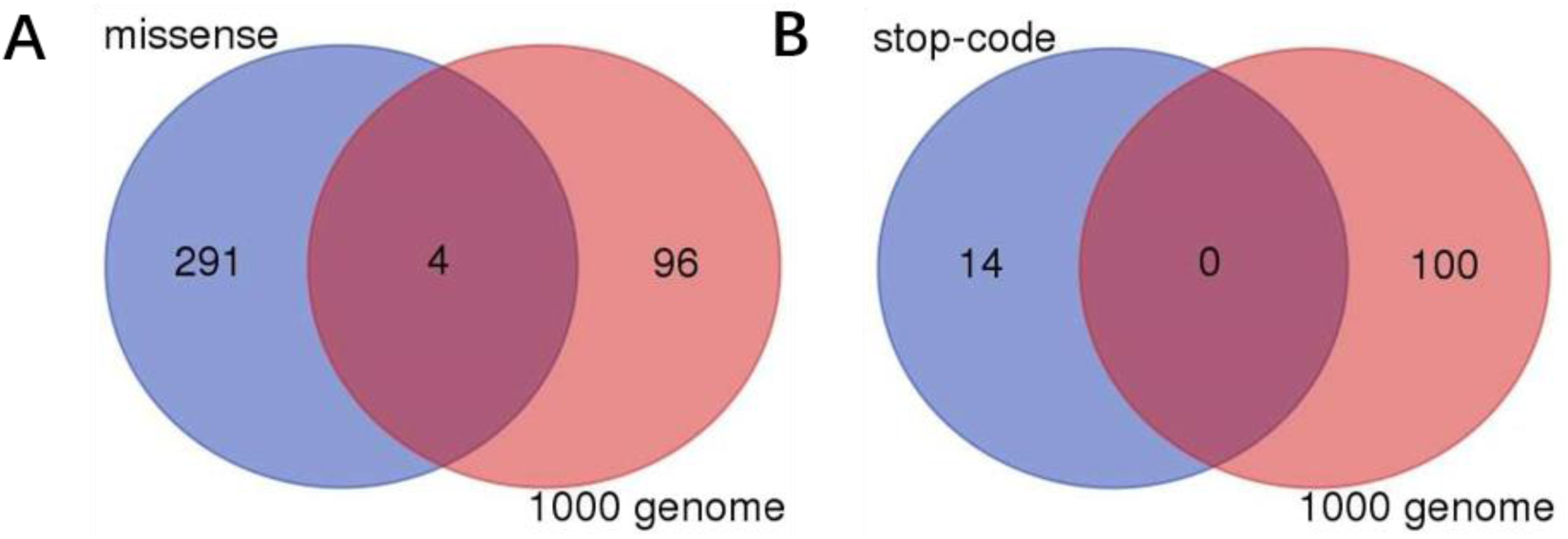
Venn diagram of the SNP screen in two database. A. the screen of the missense SNPs in NCBI database with SNPs in 1000 genome database; B. the screen of the stop-code SNPs in NCBI and SNPs in 1000 genome.

**Figure 2.**
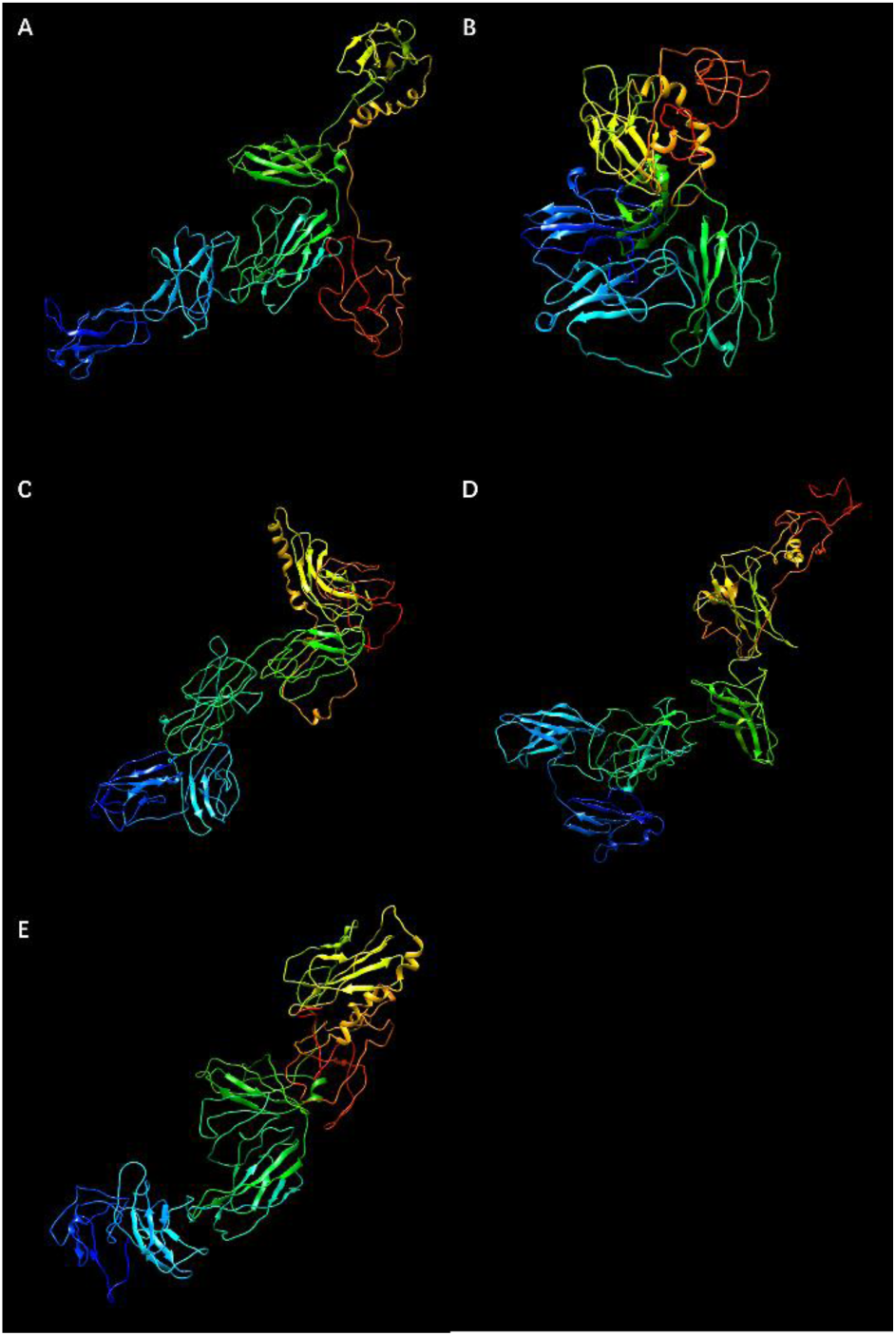
protein structure simulation with Phyre 2.0. A-E : wild type, rs190983971, rs563996763, rs546510242 and rs373621350 Respectively

According to the Uniprot database, rs190983971 linked to a ACT to GCT transition which means a T to A transition in amino acid level. Also, its mutation site, 374 from C terminal, located in the cellular immunoglobulin-like domain. Since the better hydrophily property T hold compared to A, we deduct a more stable structure the T374A holds. As shown in Figure 3 A, we can see a condenser conformation. With a condenser structure, we estimate the T374A CMPL protein will form a tighter dimer in the cell membrane and once bind to TPO, will then start a ankylosing activation on the following genes.

**Figure 3.**
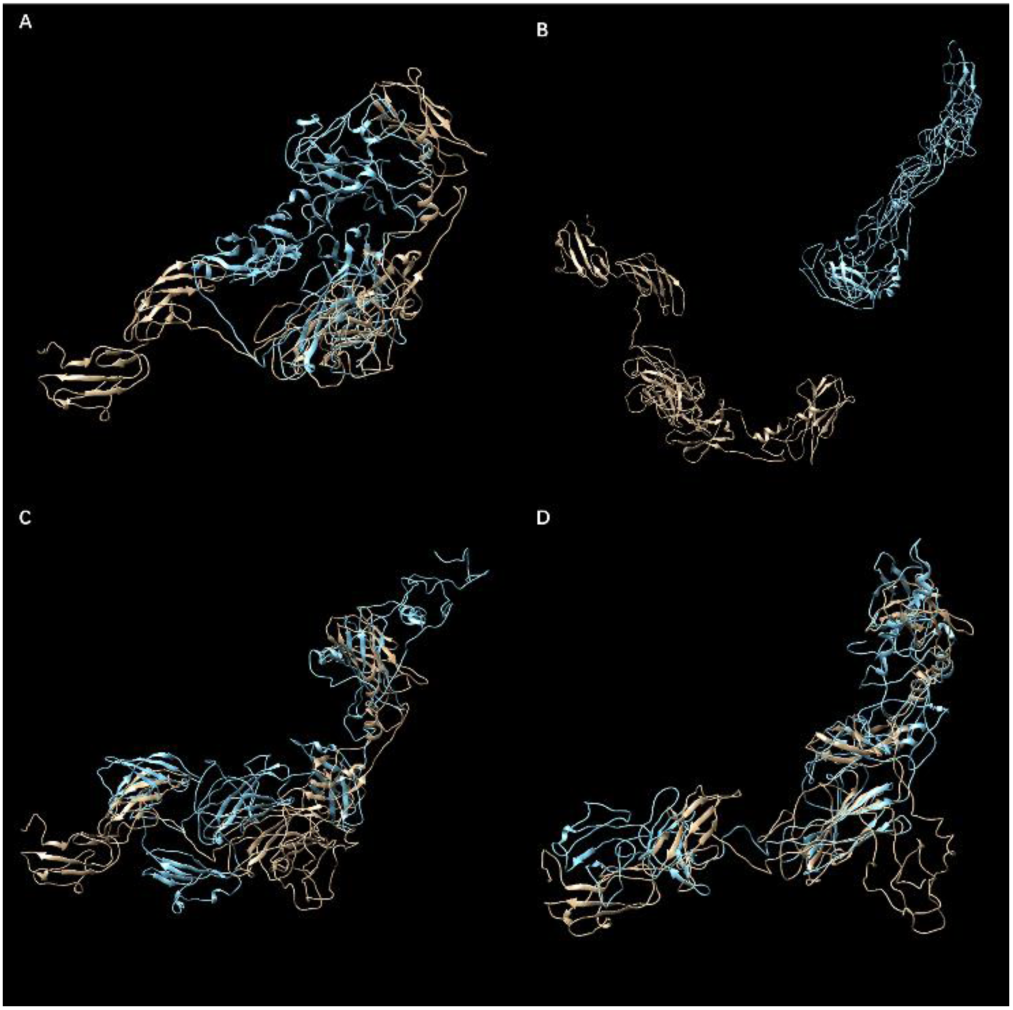
Structure comparison between wild type and mutated protein. Brown structures show wild type conformation and the blue one means mutated structure. A-D: rs190983971, rs563996763, rs546510242 and rs373621350 Respectively

In Figure 3 B, the completely divergence of the wild type and T428I (rs373621350) structure indicates the huge transformation in the secondary and tertiary structure. Located in the fibronectin-III domain, the mutation of 428 switch the D2-fragment towards the cell membrane and cause the misplacement of the D1-fragment which is connected to D2. As we shown in Figure 2 C, the extracellular region, transmembrane region and then the intracellular region are getting closer, which means the slack of their binding function to TPO.

The K355E (rs546510242) and R390C (rs373621350), when compared with wild type, shown very similar structure and hint that the amino acid substitute bring little interference to the function. (Figure 3 C, Figure 3 D) In fact, though K and E are both hydrophilic, K belongs to alkaline amino acid while E belongs to acidic amino acid. The site it place, the D1 domain, is responsible for the recognition of TPO peptide, so if 355 is the key fragment of TPO-MPL recognition, then the affinity between then may decrease.

In a word, these 4 SNPs all cause the substitution of amino acid and among then, rs190983971 and rs373621350 cause tremendous conformational changes. We believe these 4 SNPs will more or less affect the binding of MPL and TPO and then influence the signal pathway downstream.

### SNP distribution

To predict the risk of suffering Megakaryocyte Associated Diseases, we ranked the 4 SNPs and their geographical distribution in 1000 genome (Figure 4), and draw a SNP-branches Map. (Figure 5) We can then observed a enrichment of rs190983971 SNPs in BEB, CDX, CHB, CHS, JPT, KHV, PJL and STU, suggesting the high diversity and wide spreading of MPL T374A. At the same time, both rs373621350 and rs563996763 are specific to only one population, BEB and IBS respectively. The distribution region of rs190983971 is commonly seen in Eastern Asia while rs190983971 and rs546510242 are cross-link in Middle East. Interestingly, Indian peninsula congregates 3 of the SNPs we detected with rs190983971 and rs373621350 in Bangladesh and rs546510242 in India. From an Intercontinental prospective, South America, Africa, and most area of Europe show no spread of these 4 SNPs, indicating a ethnical difference in genome.

**Figure 4.**
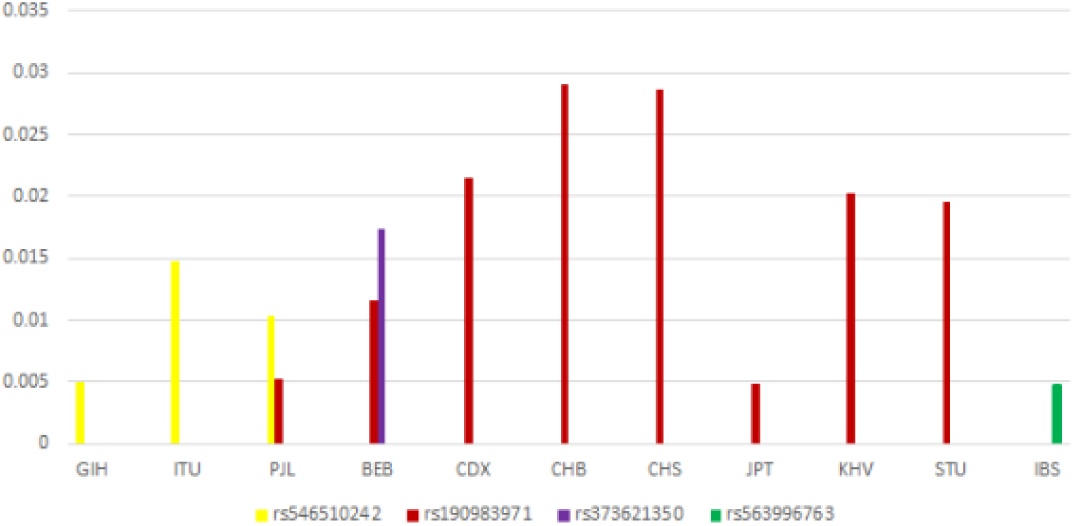
statistic Bar chart of SNPs ratio in different branches.

**Figure 5.**
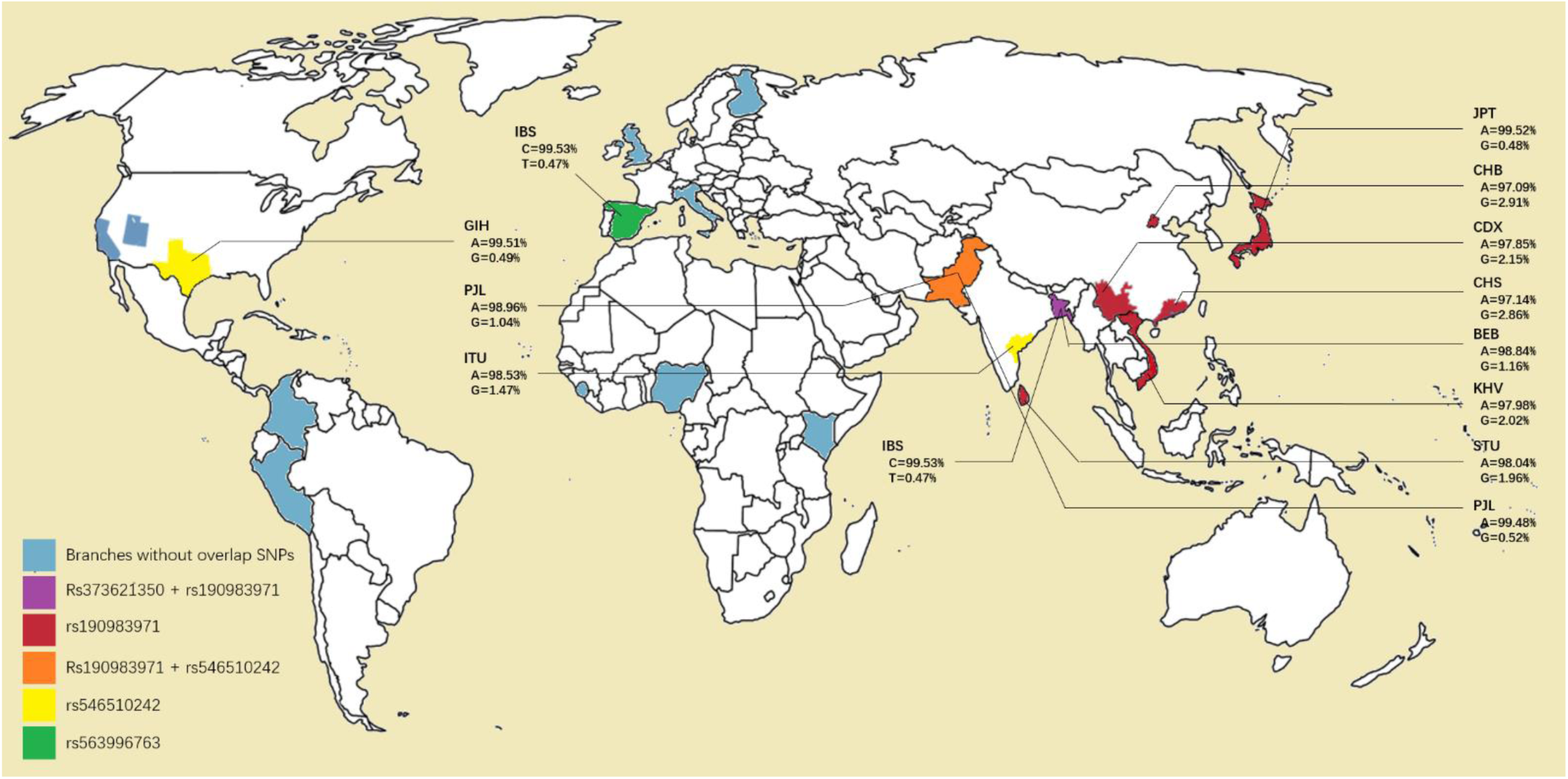
geographical distribution of *mpl* SNPs: Blue region labeled area and habitant carried no screened SNPs; Purple means branches carries rs37362150 and rs190983971 at the same time. Red region means branches carry rs19083971; Orange region manifests area where rs19083971 and rs546510242 are detected at the same time; Yellow region means branches with rs546510242 distribution; Green region means branches carrying rs563996763.

## IV. DISCUSSION

From the result of protein structure simulation, we can observe great transformation in protein structure in rs190983971 and rs563996763. However, on the one hand, in silica simulation can’t fully simulate the in vivo state and the existence cell membrane can keep the C terminal and N terminal of MPL far in two opposite side, so a more extend structure can be well kept in cells. On the other hand, rs546510242 and rs373621350, may have less conformational change, but the replacement of amino acid with different character can also lead to the increase or decrease of the affinity between MPL and TPO. Based on this work, we try to shed a light for the target of MPL gene screen and also hint the way of further research in wet lab experiment to confirm the effect of these SNPs.[7]

## REFERENCES

[1]. Hussein, K., et al., MPLW515L mutation in acute megakaryoblastic leukaemia. LEUKEMIA, 2009. 23(5): p. 852–855.

[2]. Pikman, Y., et al., MPLW515L is anovel somatic activating mutation in myelofibrosis with myeloid metaplasia. PLOS MEDICINE, 2006. 3(E2707): p. 1140–1151.

[3]. Beer, P.A., et al., MPL mutations in myeloproliferative disorders: analysis of the PT-1 cohort. BLOOD, 2008. 112(1): p. 141–149.

[4]. Loots, G.G., et al., Identification of a coordinate regulator of interleukins 4, 13, and 5 by cross-species sequence comparisons. SCIENCE, 2000. 288(5463): p. 136–140.

[5]. Gottgens, B., Analysis of vertebrate SCL loci identifies conserved enhancers (vol 18, pg 502, 2000). NATURE BIOTECHNOLOGY, 2000. 18(10): p. 1021–1021.

[6]. Tao, H., D.R. Cox and K.A. Frazer, Allele-specific KRT1 expression is a complex trait. PLOS GENETICS, 2006. 2(e936): p. 848–858.

[7]. Brix, N., et al., Arthritis as presenting manifestation of acute lymphoblastic leukaemia in children. ARCHIVES OF DISEASE IN CHILDHOOD, 2015. 100(9): p. 821–825.

